# The secreted staphylococcal biofilm protein Sbp forms biomolecular condensates in the presence of DNA

**DOI:** 10.1101/2025.10.30.685619

**Authors:** P. Ethan Adkins, Alexander E. Yarawsky, Andrew B. Herr

**Affiliations:** Division of Immunobiology, Cincinnati Children’s Hospital Medical Center, Cincinnati, Ohio, USA; Medical Scientist Training Program, University of Cincinnati College of Medicine and Cincinnati Children’s Hospital Medical Center, Cincinnati, Ohio, USA; Division of Infectious Diseases, Cincinnati Children’s Hospital Medical Center, Cincinnati, Ohio, USA; Department of Pediatrics, University of Cincinnati College of Medicine, Cincinnati, Ohio, USA

**Keywords:** Biomolecular condensate, liquid-liquid phase separation, aggregation, Small basic protein, *Staphylococcus epidermidis*, biofilm, extracellular DNA, fluorescence recovery after photobleaching, analytical ultracentrifugation, confocal microscopy

## Abstract

*Staphylococcus epidermidis* is the leading cause of device-related infections, primarily due to its ability to form biofilms, surface-adherent bacterial communities that confer remarkable resistance to antibiotics and host defenses. Small basic protein, Sbp, is a 16- kDa protein expressed by *S. epidermidis* that has been shown to be crucial for biofilm formation, but little is known about its function. Sbp features a high proportion of basic residues as well as several predicted regions of intrinsic disorder. Consistent with its high positive charge density, Sbp is shown here to interact with double-stranded DNA, a ubiquitous component of the biofilm matrix, forming soluble complexes or large aggregates with short or longer DNA oligonucleotides, respectively. The observed multivalent interaction of Sbp with DNA along with its predicted disorder suggested that it might form biomolecular condensates with DNA. Confocal and differential interference contrast microscopy revealed that Sbp and dsDNA form phase-separated droplets and/or solid aggregates, depending on the concentration and stoichiometry of Sbp and DNA as well as the DNA oligonucleotide length. Fluorescence recovery after photobleaching experiments demonstrated that Sbp-DNA condensate droplets initially exhibit liquid-like behavior but gradually transition to a gel-like state. This work provides the first evidence that Sbp binds DNA and undergoes biomolecular condensation, revealing a previously unrecognized mechanism that may contribute to biofilm matrix organization in *S. epidermidis*.

## Introduction

*Staphylococcus epidermidis* is a gram-positive commensal bacterium found ubiquitously on the skin that is the leading cause of hospital-acquired infections in the US^1^. Its success as a pathogen is attributed to its ability to form biofilms, multicellular communities surrounded by an extracellular matrix that adhere to abiotic or biotic surfaces^2; 3^. Bacteria growing in biofilms, whether found on indwelling catheters, prosthetic joints, or other medical devices, are highly tolerant to antibiotics and host defenses, making *S. epidermidis* one of the most important causes of device-related infections. The biofilm matrix is composed of several constituents, including polysaccharides, proteins, and extracellular DNA (eDNA). Recent studies have highlighted that eDNA plays an important structural role in biofilm architecture and stability^4^.

Among proteins implicated in the formation of the *S. epidermidis* biofilm matrix, the 16-kDa Small basic protein (Sbp) has emerged as a key component. Sbp was isolated from crude biofilm mixtures and shown to be necessary for robust surface colonization^5^. Sbp- deficient mutants have reduced ability to adhere to a surface and form multilayered biofilms. While Sbp was shown to localize on the bacterial cell surface, strikingly, it was also found to concentrate in a layer at the biofilm–surface interface within unevenly distributed clusters. These experiments suggest that Sbp may be involved in the development of functional scaffolding of biofilm matrix components, which, when removed, compromises biofilm development on clinically relevant surfaces.

Sbp was first identified and isolated from the biofilm matrix by affinity chromatography using Sepharose beads coupled to B-repeat domains of the accumulation-associated protein (Aap)^5^. In addition to its interaction with the B-repeats of Aap, further study showed that Sbp influenced biosynthesis of the biofilm polysaccharide PIA^5^. This evidence supports a model in which Sbp acts as a multifunctional structural adapter that promotes Aap-mediated cell–cell attachment as well as PIA-dependent processes, integrating multiple matrix components into a cohesive scaffold. This interplay with multiple matrix constituents places Sbp at an intersection between adhesion, accumulation, and matrix organization, indicating it is an important determinant of biofilm mechanics.

Double-stranded eDNA is another ubiquitous constituent of biofilm matrices. It promotes initial adhesion, acts as a structural tether between cells, chelates cations, and can nucleate assembly of other matrix components^6^. The majority of matrix eDNA in *S. epidermidis* biofilms consists of chromosomal DNA that is released from a subset of *S. epidermidis* cells by the action of autolysin E (AtlE), a lytic enzyme that breaks down the peptidoglycan components of the bacterial cell wall^7^. *S. epidermidis* AtlE-mutant strains that fail to release eDNA demonstrated inhibited ability to form biofilm matrix on polystyrene surfaces, with some mutants unable to form multilayered cell clusters^8^. The addition of DNase similarly reduced attachment of *S. epidermidis*. This data suggests that it is necessary for *S. epidermidis* to sacrifice a portion of cells to release necessary eDNA through the autolytic activity of AtlE. Importantly, several bacterial amyloid systems are promoted or stabilized by the presence of DNA. In *Staphylococcus aureus*, for example, eDNA facilitates assembly of phenol-soluble modulin amyloid fibers, and more generally eDNA–protein interactions are increasingly recognized as drivers of matrix architecture^9; 10^.

A rapidly evolving framework for understanding how macromolecules interact is the formation of biomolecular condensates, a subset of which is referred to as liquid–liquid phase separation (LLPS). Over the past decade, phase separation has been recognized as a mechanism by which proteins, nucleic acids, and their complexes demix from the surrounding solution to form biomolecular condensates with distinct composition and physical properties^11–16^. It has been observed that biomolecular condensates selectively allow or preclude diffusion of external macromolecules^17^. The mechanisms behind condensates’ abilities to limit diffusion across their interface are not fully understood.

Nucleic acids can actively promote or regulate LLPS by providing binding sites that nucleate protein assembly; conversely, protein condensation can localize nucleic acids into concentrated compartments. While LLPS has been mostly studied in eukaryotic cells, evidence indicates that phase separation also occurs in bacterial and extracellular contexts, and that it can nucleate transitions to solid states such as aggregation or amyloid fibrils^18–22^.

Several characteristics of Sbp described below suggest that it is likely to interact with double-stranded DNA (dsDNA) and a strong candidate to undergo phase separation. If Sbp forms DNA-driven condensates that subsequently mature into higher-order structures, that pathway would provide a mechanism for Sbp’s role in creating a mechanically robust and adhesive matrix. In addition, establishing Sbp–DNA phase behavior would extend LLPS concepts to extracellular matrix assembly in bacteria and suggest new avenues for targeting biofilm mechanics therapeutically.

In this report we test the central hypothesis that Sbp undergoes phase separation in the presence of double-stranded DNA, and that DNA-driven condensates provide a nucleation platform for Sbp’s transition to higher-order solid-phase assembly. Using a combination of microscopic imaging, spectroscopy, and biophysical characterization, we have demonstrated Sbp interacting with dsDNA, and mapped the conditions (protein:DNA stoichiometry, DNA length) that promote condensation and solid-phase transition. By integrating mechanistic biophysics with bacterial cell biology, our results aim to connect molecular-level phase behavior to mesoscale biofilm architecture and to suggest new molecular targets for disrupting clinically problematic *S. epidermidis* biofilms.

## Results

### Sbp has high positive charge density and predicted disordered regions

Despite its relatively small size of 16 kDa, Sbp’s primary sequence contains a high proportion of positively charged residues (Fig. 1A). Analysis using the CIDER webserver, which predicts net charge per residue (NCPR), revealed several clusters of positive charges along the sequence (Fig. 1B, 1E). Given this high charge density and the absence of experimental structural data, we used a suite of predictive tools to assess Sbp’s structural properties. Multiple predictors of intrinsic disorder consistently identified the N-terminal 10-20 residues as disordered, as well as other potentially disordered regions (Fig. 1C, 1F). Alphafold3 modeling further suggested that Sbp adopts a fold composed of a continuous antiparallel β-sheet on one side of the protein, with the N-terminus folding back across the face of the β-sheet (Fig. 1D). Although Alphafold3 predicts a compact globular fold, the high positive charge density and predicted disordered regions suggest that Sbp may instead form a less compact conformation with regions such as the N-terminus extending away from the remainder of the partially-folded protein. Furthermore, SAXS and NMR data have been reported suggesting that Sbp is partially folded^23^. For all following experiments, Sbp was recombinantly expressed as a fusion protein with an N-terminal 6xHis–maltose-binding protein (MBP) tag. The tag was removed by tobacco etch virus (TEV) protease cleavage and further purification, yielding the native Sbp sequence containing only an additional N-terminal glycine after the TEV protease cleavage.

**Figure 1.**
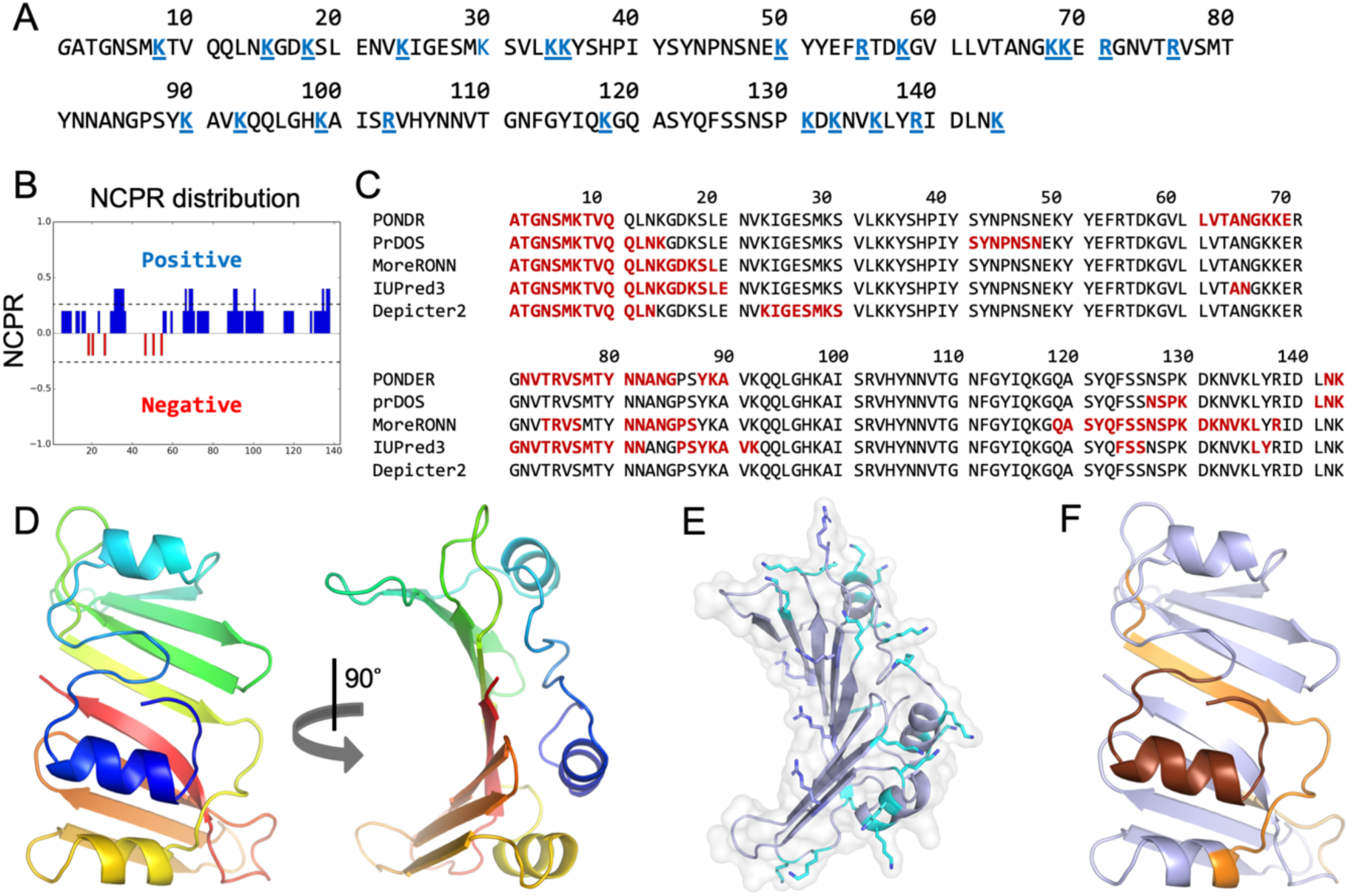
Sbp contains a high proportion of basic residues and predicted disorder. (A) The primary sequence of the Sbp construct is shown with basic residues highlighted in blue, demonstrating even distribution of positive charges throughout. This is further illustrated by panel (B), a blob index of net charge per residue (NCPR) generated by the CIDER server^54^. Panel (C) shows the results of Sbp’s primary sequence when put through several disordered region predictive models^54–57^, consistently indicating a disordered N-terminal region. (D) Alphafold-predicted structure of Sbp, colored from blue (N-terminus) to red (C-terminus). (E) Sbp model showing basic residues as colored sticks (blue, arginine; cyan, lysine). (F) Sbp model showing consensus regions predicted to be disordered as shown in panel B. Colored structural elements highlight sequence regions predicted to be disordered in at least 3 predictions: brown for N-terminal residues 1-19; orange for residues 72-90; and yellow for residues 124-130. Note that the AlphaFold models assume a fully folded protein, although the positive charge density and predicted disorder suggest that Sbp may not form a compact globular fold under physiological conditions.

### Sbp associates with double-stranded DNA and forms turbid assemblies

Sbp contains a high proportion of positively charged residues (Fig. 1) and it is known to bind to the B-repeat domain of Aap—an elongated, negatively charged protein found in *Staphylococcus epidermidis* biofilms^5^. Thus, we examined whether Sbp could also interact with negatively charged dsDNA. To test this, we used synthetic dsDNA oligomers of varying lengths composed of repeating (ACTG)n sequences. This sequence was selected as a representative oligonucleotide without sequence specificity. Oligomers are referred to as “ds–” followed by their base-pair length (e.g., ds8 is an 8 bp duplex with the sequence 5′– ACTGACTG–3′).

Sedimentation velocity analytical ultracentrifugation (SV-AUC) on Sbp alone revealed a monomeric species that sedimented at 1.58 S with a frictional ratio (f/f0) of 1.45, corresponding to a calculated molar mass of 15.9 kDa (compared to an expected molar mass of 16.2 kDa). The frictional ratio of 1.45 indicates a slightly asymmetrical shape (f/f0 of 1.0 corresponds to a perfect sphere; 1.3 – 1.4 to a mostly globular protein like bovine serum albumin; 1.6 – 1.7 to a more asymmetric protein like IgG; and >1.7 is indicative of elongated or non-globular species)^24–26^. Sbp with increasing amounts of labeled dsDNA were also analyzed by SV-AUC, revealing a shift to higher sedimentation coefficients beyond that of either Sbp or ds14 (Fig. 2A), indicating association of Sbp and ds14. The labeled dsDNA was specifically detected at 665 nm, allowing the analysis to specifically report on the sedimentation behavior of the DNA. The concentration of DNA determined by the analysis (i.e., c(*s*) peak area) was lower than expected, suggesting that significant amounts of DNA had undergone major aggregation when Sbp was present. Indeed, SV-AUC experiments using longer dsDNA oligonucleotides consistently showed a lack of observable signal, suggesting the formation of large aggregates which completely sedimented before reaching the target rotor speed.

**Figure 2.**
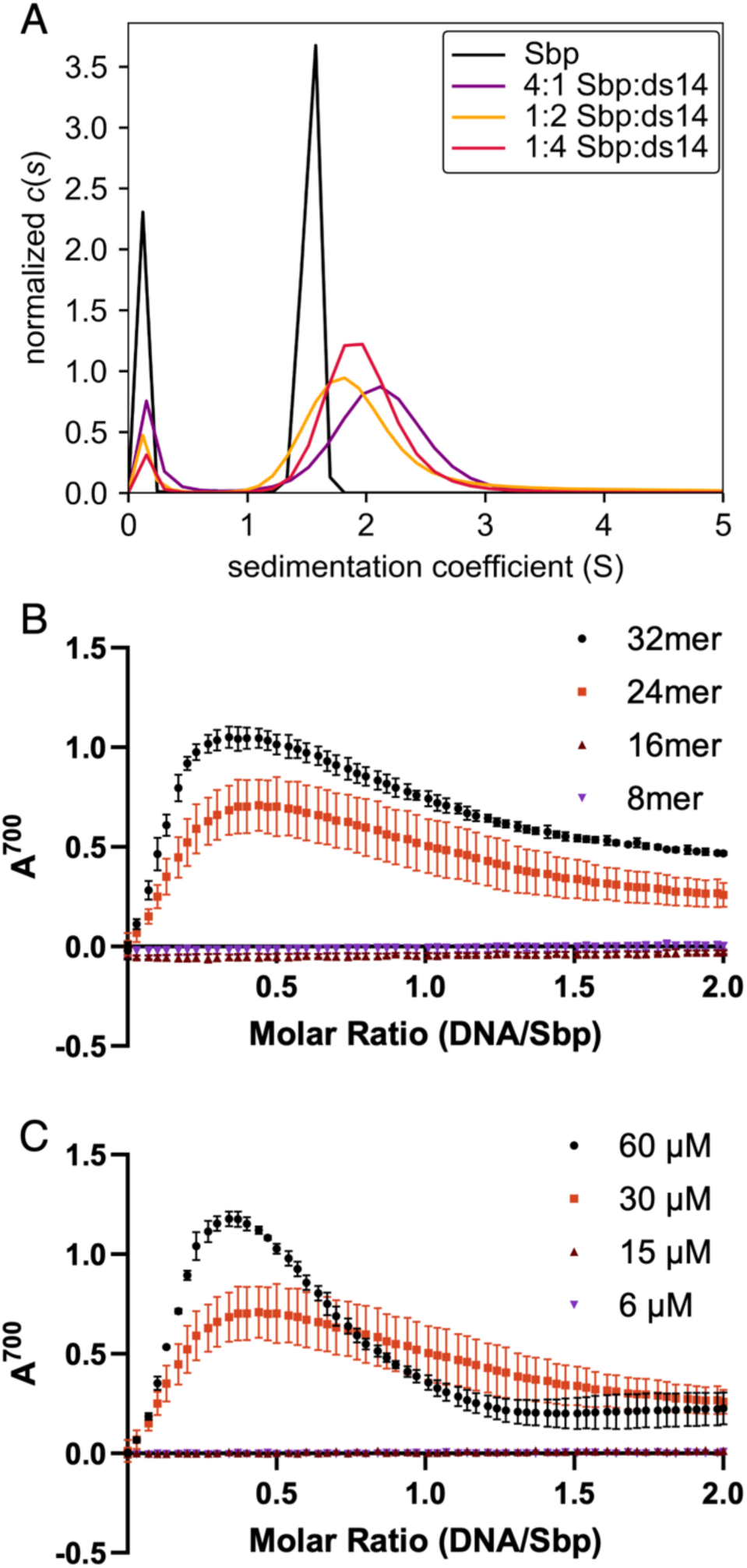
Sbp associates with double-stranded DNA, forming turbid species in the presence of dsDNA of sufficient length. (A) Sbp’s ability to associate with dsDNA was evaluated by sedimentation velocity analytical ultracentrifugation using a 14 bp dsDNA sequence (ds14). Sbp alone sediments as a momomer (black curve) whereas samples containing Sbp and ds14 shift to higher sedimentation coefficients, indicating that Sbp binds ds14 and forms stable complexes. (B) Turbidity measurements of 30 μM Sbp in the presence of increasing molar ratios of dsDNA oligomers of 8, 16, 24, and 32 bp in length indicate the formation of very large aggregates or condensates. (C) Turbidity measurements with varying starting concentrations of Sbp and increasing molar ratios of 24 bp oligomer reveal that Sbp concentrations of 30 μΜ or higher are required for formation of the large aggregates or condensates.

To better characterize the interaction of Sbp with longer dsDNA oligonucleotides, turbidity assays were carried out to measure light scattering at 700 nm after successive aliquots of ds8, ds16, ds24, or ds32 were added to 30 µM Sbp. The Sbp:dsDNA mixtures using ds8 or ds16 showed only baseline light scattering; however, addition of ds24 or ds32 showed a large increase in light scattering with increasing dsDNA, reaching maximal intensity at DNA:Sbp ratios of 0.44 and 0.34, respectively (Fig. 2b). Turbidity also depended on Sbp concentration: ds24 induced no signal when added to 6 or 15 µM Sbp but produced robust scattering at 30 and 60 µM. Together, these data demonstrate that Sbp binds dsDNA and that formation of turbid, higher-order assemblies requires dsDNA longer than 16 bp and Sbp concentrations above ∼15 µM.

### Sbp undergoes DNA-dependent liquid–liquid phase separation

Many phase-separating proteins share three features: intrinsic disorder, multivalent interactions, and nucleic acid binding^12; 16; 19; 27^. As mentioned, various disorder prediction servers indicate that Sbp has a high probability of intrinsic disorder (Fig. 1C, 1F), and Sbp has been shown to be partially folded^23^. Our SV-AUC and turbidity data indicate that Sbp can interact multivalently with DNA to form massive aggregates that pellet immediately in the analytical ultracentrifuge. We therefore tested whether Sbp-DNA mixtures can undergo liquid-liquid phase separation to form biomolecular condensates.

Turbidity assays suggested the formation of large assemblies when Sbp was mixed with sufficiently long dsDNA (Fig. 2). Turbidity alone cannot distinguish solid aggregation from liquid–liquid phase separation (LLPS), therefore, we performed differential interference contrast (DIC) microscopy. Within 30 s of addition of dsDNA, Sbp formed spherical, liquid-like droplets that ranged from 0.5 to 1 μm in diameter (Fig. 3A). At higher DNA:Sbp ratios, we also observed solid aggregates coexisting with droplets. Confocal imaging using Dylight488-labeled Sbp and TYE665-labeled dsDNA confirmed that both components were enriched within the dense phase (Fig. 3B). Over time, DIC imaging revealed droplet growth and coarsening, followed by the appearance and enlargement of solid aggregates (Fig. 3C, Supplementary Movie).

**Figure 3.**
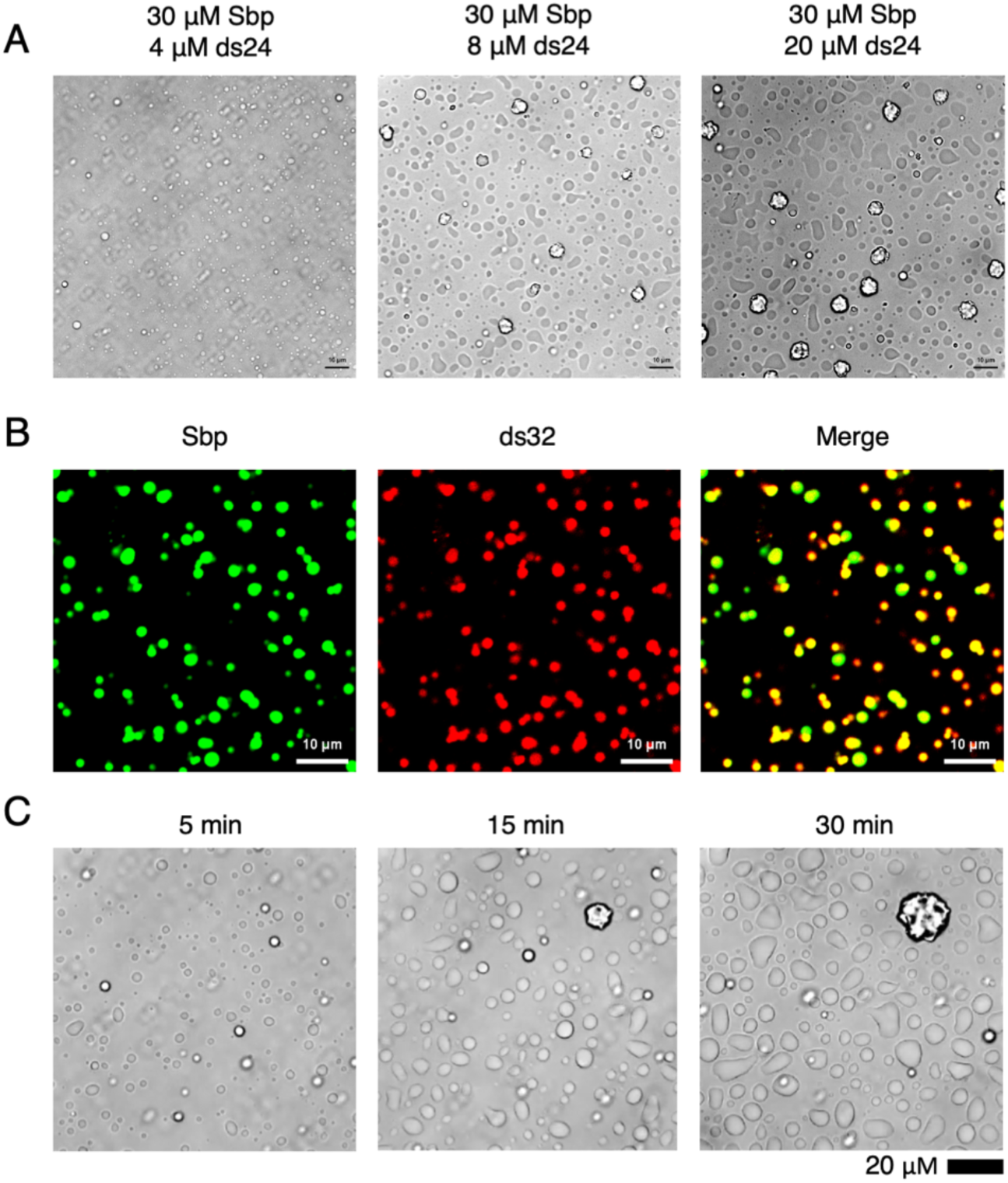
Sbp forms phase-separated droplets and solid aggregates in the presence of dsDNA in vitro. (A) Differential interference contrast (DIC) microscopic images of Sbp reveal the formation of droplets and/or aggregates after the addition of 32mer dsDNA. Higher concentrations of DNA lead to more rapid droplet/aggregate formation and growth. Confocal imaging of 60 μM Sbp (Dylight488-labeled) and 4 μM ds32 (TYE665- labeled) is shown in (B). Only liquid droplets are observed in these conditions, and the merged image shows that droplets are enriched in both Sbp and ds32. (C) Time-lapse DIC images of 30 μM Sbp and 8 μM ds32 demonstrate liquid droplet and solid aggregate growth over time.

We quantified droplet size over time using confocal microscopy with mixtures of Sbp with labeled ds24, ds32, and ds60 at 22 °C and 37 °C (Fig. 4A, 4B). Droplets increased in size from less than 1 μm up to 5 μm in diameter over a period of 30 minutes and gradually merged with other droplets to eventually form a bulk condensate dense phase. The temperature dependence of droplet size varied based on the dsDNA oligonucleotide length.

**Figure 4.**
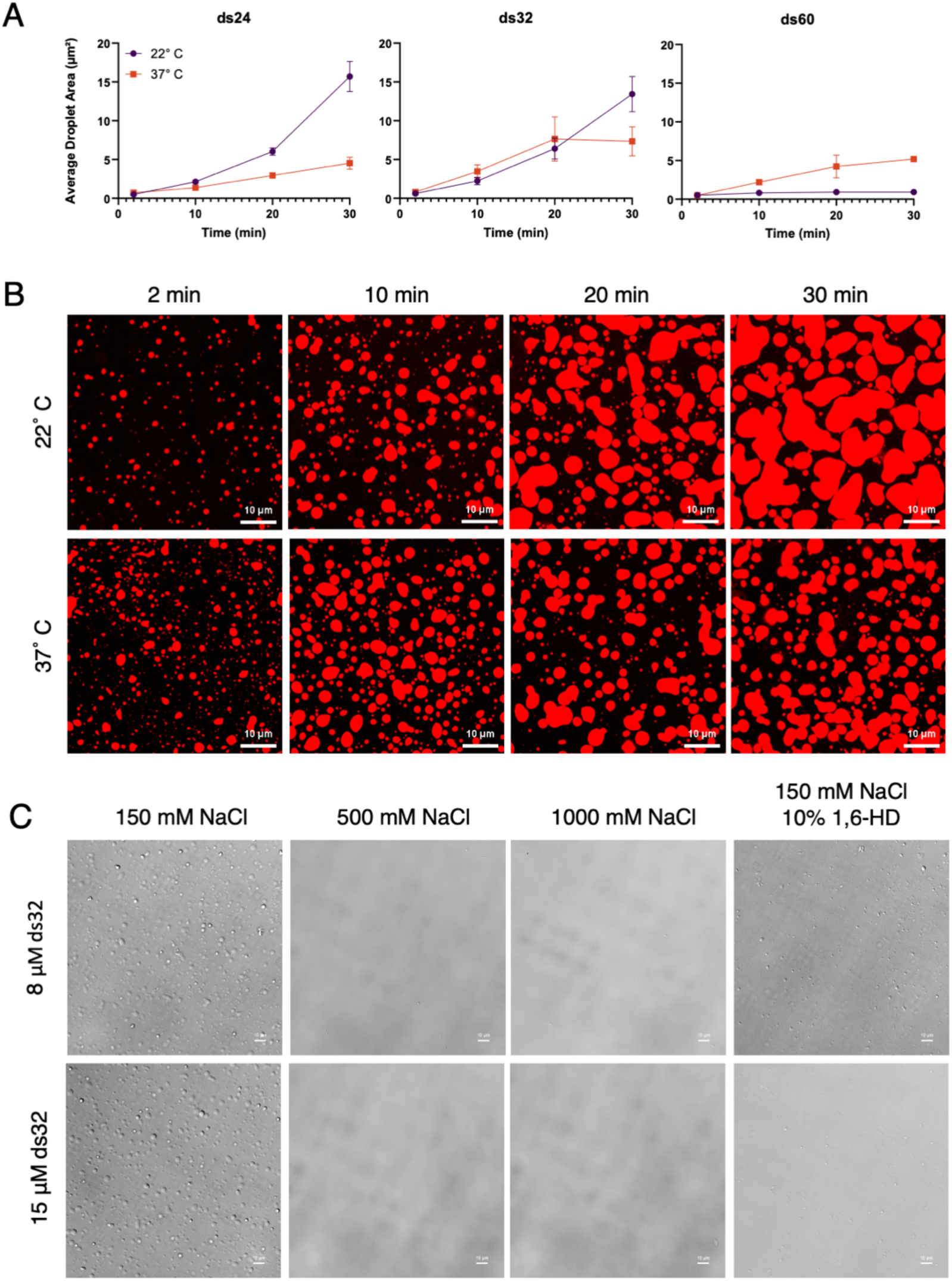
Characteristics of Sbp:dsDNA condensate droplets. (A) Average size of Sbp-ds32 droplets at progressing time points, observed at 22° and 37° C. Nikon Elements software Object Count tool was used with smoothing and separation parameters adjusted to count merged droplets as multiple individual droplets. (B) Representative micrographs from experiments in Fig.6A. (C) DIC images of 45 μM Sbp and 8/15 μM ds32 with increased concentration of buffer NaCl or addition of 10% 1,6- hexanediol. DNA concentrations were chosen as conditions where only LLPS (8 μM) or LLPS+Aggregation (15 μM) were present, respectively. 500 and 1000 mM NaCl conditions completely abrogated phase separation. 10% 1,6-hexanediol severely inhibited, but did not prevent, condensate formation.

### Both hydrophobic and electrostatic interactions drive condensate formation

There are many types of interactions that promote formation of condensates^28; 29^, but some of the most frequently observed are hydrophobic interactions (commonly involving aromatic residues), cation-π interactions (especially involving Arg residues), and charge-charge interactions^28; 30–32^. We varied solution conditions to provide insight into the interactions that stabilize Sbp:dsDNA condensates. The aliphatic alcohol 1,6-hexanediol can often inhibit LLPS by disrupting hydrophobic protein-protein interactions but does not affect charge-mediated condensates^33^. On the other hand, charge-based interactions between proteins and nucleic acids are inhibited at high NaCl concentrations (approaching 1 M), since high concentrations of counterions effectively screen electrostatic interactions to the extent that they are negligible at 1 M NaCl^34^. Using DIC microscopy, we observed that 500 mM or 1 M NaCl completely inhibited formation of both condensate droplets and aggregates formed by 45 μM Sbp and either 8 or 15 ds32 (chosen to optimize formation of only droplets versus droplets with aggregates, respectively). Addition of 10% 1,6- hexanediol greatly inhibited both droplets and aggregates but did not completely block their formation (Fig. 4C). The observed NaCl dependence highlights the importance of electrostatic interactions between Sbp and dsDNA in driving both phase separation and aggregation in this system. However, the inhibitory action of 1,6-hexanediol reveals that hydrophobic interactions are also involved in condensate and aggregate formation by Sbp and dsDNA.

### Phase diagrams for formation of Sbp:dsDNA droplets and solid-phase aggregates

We next mapped the conditions under which Sbp and dsDNA form droplets, aggregates, or both. Using DIC microscopy, we constructed phase diagrams by mixing Sbp with dsDNA oligomers at varying molar ratios, gently stirring, and imaging after 5 min (Fig. 5a). Although mixtures of Sbp with ds16 did not induce detectable turbidity as measured by light scattering, DIC revealed small droplets at a minimum of 30 μM Sbp, indicating that dsDNA shorter than 24 bp can nucleate LLPS under certain conditions. However, oligonucleotides ranging from ds8 through ds14 did not show any condensate formation. Droplets formed by ds16 were consistently smaller than those driven by longer oligonucleotides, and most conditions under which Sbp:ds16 formed droplets were free of aggregates. Ds24 induced aggregate-free phase separation under a range of conditions starting predominantly at 30 μM Sbp:8 μM ds24 and ranging to 92.5 μΜ Sbp with 8 – 24 μΜ ds24 (Fig. 5a). Sbp concentrations between 55 – 92.5 μΜ formed aggregate-free droplets under the widest range of ds24 concentrations (ranging from 8 - 38 μΜ dsDNA). The highest concentrations of ds24 consistently led to formation of both droplets and solid-phase aggregates. When ds32 was added to Sbp, a similar combination of conditions resulting in either pure phase separation or a mixture of droplets and aggregates was observed, but the region of aggregate-free droplets occurred predominantly at lower concentrations of Sbp.

**Figure 5.**
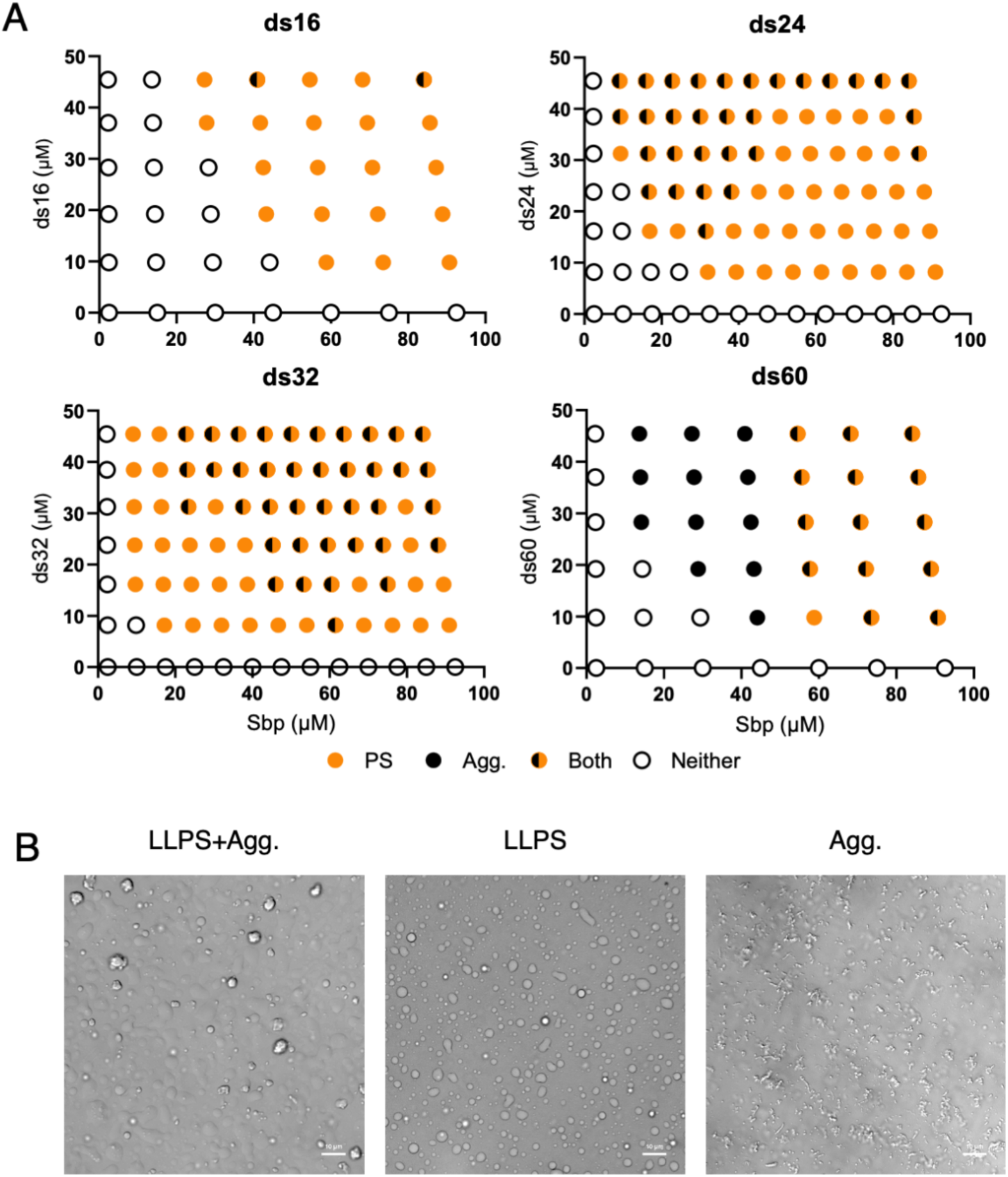
Sbp and dsDNA form liquid droplets and solid aggregates as a function of their relative molar ratios and dsDNA oligonucleotide length. Panels in (A) show diagrams of phase separation of Sbp with ds16, 24, 32, and 60. DNA was added to Sbp and allowed to incubate for 5 minutes before DIC imaging. Symbols indicate the presence of phase-separated droplets, solid-phase aggregate, both, or neither. (B) shows representative DIC images of each type of condition.

Unlike the shorter dsDNA oligonucleotides, ds60 mixed with Sbp favored pure aggregate formation (without droplets) in nearly half of all conditions, with higher Sbp concentrations favoring a mix of droplets and aggregates. The overall trend shown indicates that longer oligonucleotides require lower Sbp concentrations to phase separate, and that ds60 starts to favor aggregation over droplet formation. Representative images showing conditions with droplets only, droplets and aggregate, or aggregate only are shown in Fig. 5B.

Across nearly all tested concentrations, increasing the DNA:Sbp ratio promoted the appearance of solid aggregates alongside droplets. Time-lapse imaging showed that after initial droplet growth plateaued, droplet size began to decrease while aggregates continued to expand (Fig. 3c), suggesting a gradual recruitment of material from the liquid phase into solid assemblies.

### Sbp:dsDNA droplets initially exhibit liquid-like dynamics but mature into gel-like states

To probe the material properties of Sbp–DNA droplets, we investigated the fluorescence recovery after photobleaching (FRAP) on dense-phase droplets containing TYE665-labeled ds32. FRAP involves laser bleaching a region of interest within a droplet and measuring the recovery of fluorescence within the bleached region. The rate of fluorescence recovery in the simplest cases is determined by the rate of diffusion of the photobleached macromolecule, with a higher recovery rate occurring in more liquid-like states. However, in many cases the rate of recovery is dependent on additional factors, including the concentration of the labeled macromolecule; its rate of association with other less-labile macromolecular species; and the affinity and stoichiometry of its binding partners. Furthermore, assembly mechanisms other than LLPS such as low-valency interactions with spatially clustered binding sites^35^ can also show FRAP recovery curves that appear similar to those seen for LLPS. However, FRAP analyses can still provide useful information, such as determining the mobile fraction within the condensate droplet from the plateau of the fluorescence recovery curve (normalized to the pre-bleach fluorescence level). Condensate droplets that undergo maturation to form gel-like solids, for example, would show a decrease in both the recovery rate and the mobile fraction as they transition from liquid-like to gel-like condensates^36–38^.

After initial mixing of Sbp and ds32 (conditions chosen to form large enough droplets for FRAP analysis), droplets were allowed to sit at 22° C or 37° C for time periods ranging from 5 min to 2 hours prior to photobleaching and measuring the rate of fluorescence recovery. (Fig. 6). Droplets formed at 37° C initially showed liquid-like behavior with rapid fluorescence recovery with mobile fractions approaching 100% (Fig. 6A), but as droplets aged the recovery slowed and importantly, the mobile fraction decreased dramatically. In contrast, droplets formed at 22° C showed much lower recovery rates and smaller mobile fractions, even at early time points. We optimized the conditions to produce more reproducible FRAP recovery curves and more consistent bleach depth with droplets aged for 15, 30, and 60 min and fitted the average response of curves from 3 droplets in each condition to a single-exponential equation (Fig. 6B, 6C); these fits showed that the mobile fraction for the droplets at 37° C ranged from 88.4% down to 19.1% as the droplets aged from 15 min to 60 min. In contrast, the droplets at 22° C only had a maximum mobile fraction of 24.6% that decreased to 8.7% over the same time frame. These data indicate that the condensates formed by Sbp and dsDNA at physiological temperature mature over the course of minutes from liquid-like to gel-like behavior, as previously reported for many condensate systems. In particular, the formation of condensates by cGAS in the presence of dsDNA also showed a transition from liquid-like to gel-like behavior over a similar time range^27^.

**Figure 6.**
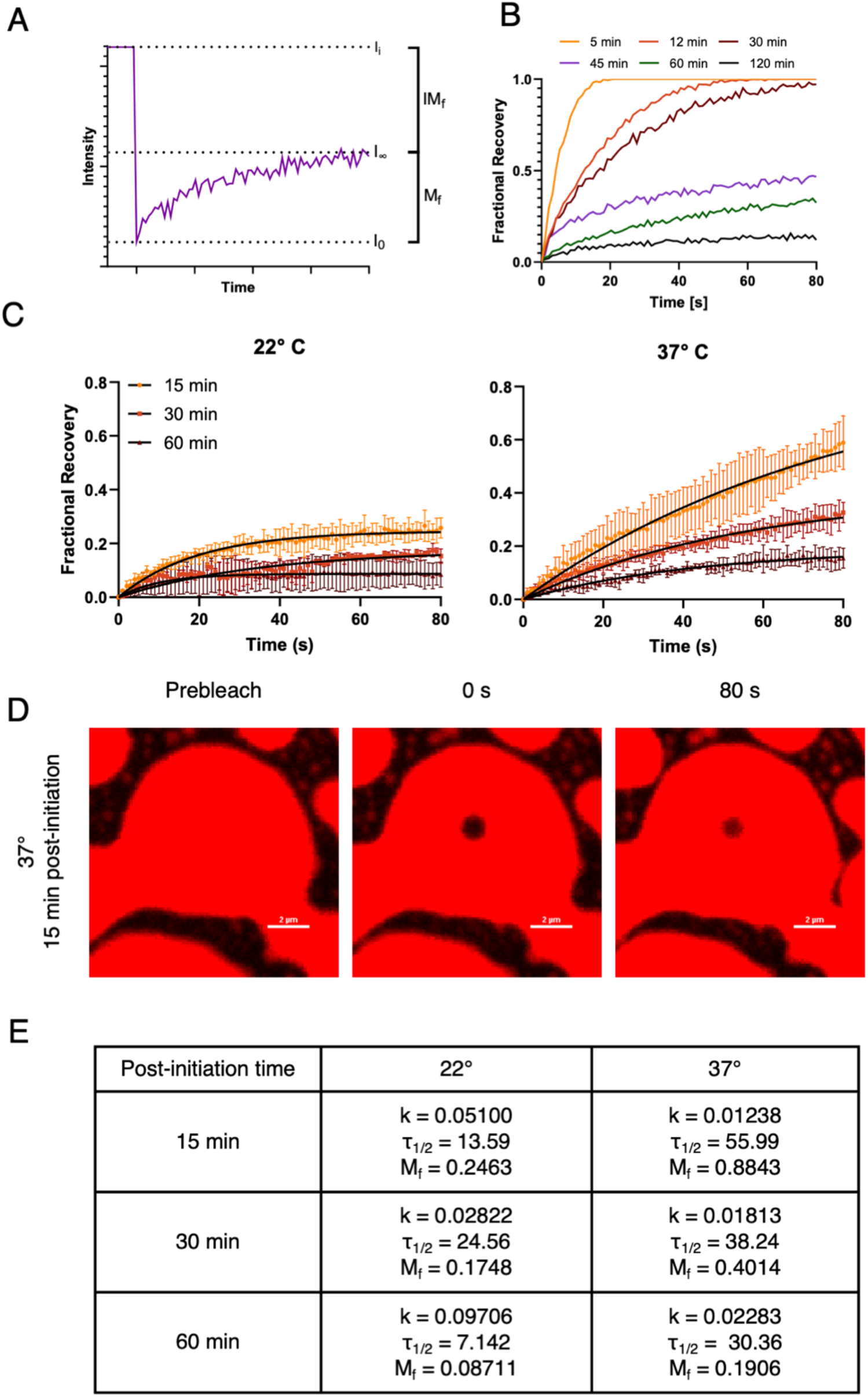
Condensate droplets formed by Sbp and DNA initially show liquid-like behavior but mature to a gel-like state. (A) A single replicate of raw FRAP data demonstrating calculated parameters. I_i_: pre- bleach fluorescence intensity, I_0_: post-bleach fluorescence intensity minimum, I_∞_: fluorescence intensity recovery plateau, IM_5_: immobile fraction, or the fraction of labeled molecules not contributing to fluorescence recovery, M_5_: mobile fraction, or the fraction of labeled molecules contributing to fluorescence recovery. (B) Normalized FRAP data qualitatively demonstrating the transition of Sbp+dsDNA condensate droplets from a liquid-like response at early time points post-mixing to a slower recovery and diminished mobile fraction as droplets mature. (C) FRAP recovery curves of Sbp and ds32 performed at 22° and 37° C after aging droplets for 15, 30, and 60 minutes. Initial time points show rapid fluorescence recovery with a mobile fraction approaching 100% but later time points progressively show slower recovery and decreased mobile fractions. Each curve represents the average of three experiments. Fits to a single-exponential model are shown in black overlaying the data points. FRAP experiments performed at 30 and 60 min were done on bulk condensate, as individual droplets were not found. (D) Representative micrographs from FRAP experiments. (E) Parameters from the single-exponential model used to fit the curves in Fig. 6A; k: exponential constant, τ1/2: half-life of fluorescence recovery, and Mf: mobile fraction.

## Discussion

Sbp is a small, secreted protein with a high proportion of positively charged residues and several regions of predicted disorder. This propensity for disorder, combined with its ability to interact multivalently with dsDNA, are common characteristics of proteins that undergo LLPS. We have demonstrated using a variety of techniques that Sbp does indeed form biomolecular condensates in the presence of dsDNA oligonucleotides of sufficient length (16 bp or longer). This dsDNA length dependence is consistent with the need for multivalency to enable condensate formation; shorter dsDNA oligonucleotides ( ≤14 bp) are presumably too short to allow the necessary degree of multivalent interactions with Sbp proteins. The findings presented here are the first to demonstrate the Sbp-DNA interaction, as well as the ability of Sbp to form biomolecular condensates.

The condensates described here form over a wide range of Sbp and dsDNA concentrations, using dsDNA oligonucleotides that range from 16 to 60 bp, and under buffer, salt, and temperature conditions that are relevant physiologically. Furthermore, it was not necessary to use crowding agents to drive condensation formation; Sbp concentrations as low as 10 μM (0.16 mg/ml) were sufficient when mixed with ds24 or ds32 oligonucleotides. The Sbp:dsDNA condensate droplets initially exhibit liquid-like behavior, including spherical morphology, the ability of droplets to fuse, and rapid FRAP recovery rates with mobile fractions near 100%. However, the droplets mature over the course of minutes to hours into gel-like or solid states. In general, although biomolecular condensates often start as liquid-liquid phase-separated droplets, the category of condensates can actually be considered a continuum that ranges from liquid-like through increasingly viscous stages to gel-like or glassy states^36^.

In the case of Sbp:dsDNA condensates, we observe a rather rapid transition from liquid-like to gel-like droplets at the same time that the droplets are merging into a bulk condensate phase. This transition between liquid and gel-like or even solid phase has been observed in many other systems^19–22^, and a common theme in many cases is that the matured droplets can act as a nucleation site for growth of amyloid fibers^19; 39; 40^. A key feature of condensate formation is the local enrichment of molecular components through dense phase coacervation. Such compartmentalization can effectively create “reaction centers,” where the elevated local concentrations of the condensate macromolecules enhance binding and reaction kinetics without requiring increased synthesis of either component. Such conditions would create an ideal local environment for assembly processes such as amyloidogenesis.

Interestingly, functional amyloid fibers form in biofilms formed by many bacterial species, where they are thought to provide resilience and physical strength to the biofilms^41–45^. In the case of staphylococci, several amyloidogenic proteins have been reported, including Aap^46–48^, phenol-soluble modulins^49^, and Sbp itself^23^. High concentrations (∼4 mg/ml) of purified Sbp have been shown to form amyloid fibrils in vitro after agitation for 12 h at 37° C^23^; it is possible that the high local concentration of Sbp in the dense phase of condensates might be sufficient to nucleate Sbp amyloidogenesis in the biofilm matrix in vivo. Furthermore, Sbp was originally identified by its ability to interact with the B-repeat superdomain from the cell wall attached protein Aap^5^. The B-repeats of Aap assemble in the presence of Zn^2+^ to form dimers and tetramers, and the tetramers nucleate functional amyloid fibers at physiological temperature^46–48; 50; 51^. Amyloid-like fibers isolated from *S. epidermidis* biofilms contained Aap as a major component, confirming that Aap amyloidogenesis occurs in vivo in staphylococcal biofilms^46^. Thus, since Sbp can bind to Aap B-repeats, Sbp-containing condensates might also act as nucleation centers for amyloidogenesis by Aap within the biofilm matrix.

It has been shown that secretion of Sbp is essential for staphylococcal biofilm formation ^5, 23^ but it was not entirely clear what role Sbp played in biofilm development. Our data suggest that Sbp acts as a central hub protein that helps to organize the scaffolding behavior of eDNA and Aap, two major macromolecular components within the staphylococcal biofilm matrix. Sbp can accomplish this function through direct interactions with both eDNA and Aap, but the ability to form condensates provides an additional level of control by allowing the potential nucleation of amyloid fibrils by Aap and/or Sbp. However, it is also tempting to speculate that Sbp:dsDNA condensates may also influence biofilm resilience by acting as selective diffusion barriers. We have observed that Sbp:dsDNA droplets merge into bulk condensate; the restricted permeability of the dense phase within such a bulk condensate in the biofilm matrix could impede the penetration of antibiotics, antimicrobial peptides, or antibodies, reducing their effectiveness.

In summary, Sbp represents a previously unrecognized example of a staphylococcal protein that undergoes DNA-dependent LLPS. Its highly charged nature enables multivalent electrostatic interactions with dsDNA, leading to condensate formation and maturation into solid precipitates. These findings broaden the understanding of how simple bacterial proteins can use phase separation to organize extracellular structures, and they highlight Sbp’s role in biofilm matrix production through coacervation with DNA.

## Materials and Methods

### Protein expression and cloning

The *sbp* gene was synthesized by IDT (Integrated DNA Technologies) with an added N-terminal Tobacco etch virus (TEV) protease cleavage site (UniProt accession no.

Q5HRC3 - aa27-169). TEV cleavage leaves an N-terminal glycine residue, which was accounted for in subsequent calculations. The IDT gBlock was inserted into the pHisMBP- DEST expression vector using the Gateway cloning system (Invitrogen). The destination vector was provided by Dr. Artem Evdokimov and contains an N-terminal 6xHis tag and maltose binding protein (MBP), which are both removed after TEV protease cleavage.

Protein was expressed in BLR(DE3) competent *E. coli* cells. One-liter cultures were inoculated with His-MBP-Sbp/BLR(DE3) and grown to an OD600 of 0.8-1.0 (approx. 4 h). Cultures were then cooled to 10° C in ice water to induce cold shock before the addition of 20 mL 100% EtOH (2% final) to maintain expression of cold chaperonins. 200 μM IPTG was used to induce expression for 16 h at 20° C. The cells were then harvested, resuspended, frozen, and thawed prior to lysis by sonication. Lysate was centrifuged, and the supernatant fraction was isolated for further purification.

### Protein purification

Sbp was purified via a 20 mL Ni^2+^ HiTrap cartridge column (GE Healthcare). The binding and wash buffer contained 20 mM Tris pH 7.4, 500 mM NaCl, 10 mM imidazole, and the protein was eluted using a linear imidazole gradient to 1 M imidazole. Eluted protein was buffer exchanged into 20 mM Tris pH 7.4, 150 mM NaCl using a 53 mL HiPrep Desalting column (GE Healthcare) before adding TEV protease in the presence of 5 mM 2- mercaptoethanol. The cleavage reaction was allowed to run at room temperature until 6xHis-MBP removal was complete. The mixture was then run over a 5 mL SP XL ion exchange column (GE Healthcare), and an NaCl gradient was used to elute Sbp.

### Turbidity Assays

For each sample, 200 μl of 30 μM Sbp was loaded into a quartz microcuvette. DNA stock solution was titrated into the sample of Sbp in 2 μL aliquots and stirred lightly using a pipette tip. After resting for 5 min to allow for any droplets or aggregate to form, the sample was stirred again and the measurement was taken. Data were recorded using a Thermo Scientific BioMate 3S UV-Visible Spectrophotometer at 280 nm to ensure accurate starting protein concentrations, as well as 400 nm and 700 nm. The data presented were measured at 700 nm, which contained data within the linear range and avoided any interference from the absorbance of protein or buffer components, while still capturing phase-separated droplets and/or aggregates. All turbidity assays were performed in 20 mM Tris, 150 mM NaCl, pH 7.4.

### Differential interference contrast microscopy

Biomolecular condensates were imaged using a Nikon TiE inverted SpectraX widefield microscope equipped with a Nikon A1 camera, with a 60x oil objective. Samples were imaged in 96 Well Square Glass Bottom µ-Plates (Ibidi). To construct phase diagrams, dsDNA was added to 150 μL of Sbp and briefly stirred. After resting for 5 min to allow phase separation to occur, images were taken. All DIC imaging for phase diagram samples were in 20 mM Tris,150 NaCl, pH 7.4, at 22° C. Samples used in DIC images in Figure 6 were in 20 mM Tris, pH 7.4, with NaCl of 150, 500, or 1000 mM (where noted).

### Confocal microscopy

Biomolecular condensates were imaged using a Nikon A1R confocal microscope with a 100x oil objective, Nikon A1 camera, and X-cite 120LED laser. Images were analyzed using Nikon Elements AR software. Nikon Elements Object Count Tool was used for average droplet area analysis, with smoothing and separation parameters adjusted. Sbp was labeled using DyLight 488 NHS Ester (Thermo Scientific). 500 μL of 1 mg/mL Sbp (60 μM) was added to a vial containing 50 μg of dye and incubated at room temperature for 1 h. Excess dye was removed through dialysis at room temperature. Degree of labeling (DOL) was calculated to be 0.4. TYE-665 labeled dsDNA was ordered directly from Integrated DNA Technologies and diluted with unlabeled DNA to a DOL of 0.5.

### FRAP assays

Fluorescence recovery after photobleaching (FRAP) assays were performed with Nikon A1 confocal microscope at 22 °C or 37 °C. TYE665-labeled ds32 was used for these experiments, as confocal imaging had established Sbp enrichment in phase separated droplets, and to avoid any alteration in the ability of Sbp to interact with dsDNA due to loss of primary amine(s) after Dylight488 labeling. Regions of interest (ROIs) with diameters of 0.5 - 1 μm were bleached using 665-nm lasers at 10% laser power for 5 seconds. Time-lapse images were acquired over an 80-second time course post-bleaching with 1-second intervals. Images were analyzed using Nikon Elements software, and normalized data was fitted using GraphPad Prism 10 to the single-exponential equation^52^:

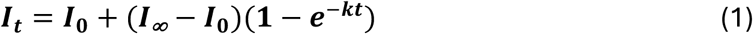

where *I0* is the lowest point of intensity (immediately post-bleach), *k* is the rate constant, and *I*∞ is the recovery plateau intensity. The mobile fraction and half-life for each FRAP recovery curve were calculated based on the equations:

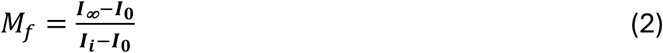

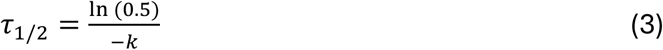

where *Mf* is the mobile fraction, *I0* and *I*∞ are as defined above, *Ii* is the intensity before bleaching, *τ1/2* is the half-life of the FRAP recovery curve, and *k* is the recovery rate fitted in equation 1.

### SV-AUC

Experiments were performed on a Beckman Coulter XL-I analytical ultracentrifuge with absorbance and interference optics as previously described^46; 47; 53^. All experiments were done at 48,000 rpm and 20° C with absorbance detection at 280 nm. Two-sector epon-charcoal centerpieces and sapphire windows were used. 440 μL of sample and matching buffer were loaded into the respective chambers and sedimentation velocity experiments were run overnight to allow for full sedimentation. SEDFIT’s continuous c(s) distribution model was used for analysis of data. All sedimentation velocity experiments were done in 20 mM Tris, 150 mM NaCl, pH 7.4, with 45 μM Sbp and 0, 11.25, 90, or 180 μM ds14. Reported sedimentation coefficients are apparent and not corrected for standard conditions.

### Supplementary material description

A supplementary movie file shows a DIC microscopy time course of 32.5 μM Sbp with 8 μM ds32 sped up 120X to illustrate the formation of condensate droplets and aggregates and the eventual merging of droplets into bulk condensate.

Filename: Adkins_Supp_Movie.MP4

## Acknowledgments

The authors gratefully acknowledge support for this work from NIGMS under R35 GM151986 (to ABH) and T32 GM063483 as well as an anonymous MSTP donor (to PEA). We thank Drs. Carlos Castañeda and Nandan Gokhale for helpful discussions and comments on the manuscript.

## Abbreviations and symbols

eDNA: extracellular DNA
Sbp: small basic protein
Aap: accumulation-associated protein
PIA: polysaccharide intercellular adhesin
AtlE: autolysin E
LLPS: liquid-liquid phase separation
dsDNA: double-stranded
DNA NCPR: net charge per residue
MBP: maltose-binding protein
TEV: tobacco etch virus
ACTG: adenine cytosine guanine,
SV-AUC: sedimentation velocity analytical ultracentrifugation
S: svedbergs
f: frictional coefficient
f0: frictional coefficient of an ideal sphere of equivalent volume
DIC: differential interference contrast
FRAP: fluorescence recovery after photobleaching
cGAS: cyclic GMP-AMP synthase
*I*∞: FRAP recovery plateau intensity
*Mf*: mobile fraction
*τ1/2*: half-life of the FRAP recovery curve

